# *Dickeya ananae* sp. nov., pectinolytic bacterium isolated from pineapple (*Ananas comosus*)

**DOI:** 10.1101/2024.10.29.620964

**Authors:** Shefali Dobhal, Nicole Hugouvieux-Cotte-Pattat, Dario Arizala, Gamze Boluk Sari, Shu-Cheng Chuang, Anne M. Alvarez, Mohammad Arif

## Abstract

Recently, species clustering within *Dickeya zeae* has been identified as complex, encompassing validly published names, including *D. oryzae* and *D. parazeae*, with some strains potentially delineating new species. In this study, genomes of strains isolated from a bacterial heart rot outbreak in pineapple (*Ananas comosus* var. *comosus*) on Oahu, Hawaii, along with two strains from pineapple in Malaysia, were sequenced. Orthologous average nucleotide identity (ANI) and digital DNA-DNA hybridization (dDDH) values among the sequenced genomes ranged from 98.93-99.9% and 91.8-99.9%, respectively, supporting the classification of seven strains within the same species. Comparisons of ANI and dDDH values between these seven strains and type strains of *D. zeae, D. parazeae,* and *D. oryzae* ranged from 94.4-95.9% and 57.2-66.5%, respectively. These values fall below the proposed boundaries for new species designation, supporting the delineation of a novel species. Phylogenetic analyses, including 16S rRNA, *gapA*, multi-locus sequence analysis (MLSA) of 10 housekeeping genes, whole-genome, and pangenome analyses, were concordant and revealed a distinct monophyletic clade, separating these strains from other members of the *D. zeae* complex, with *D. oryzae* as the closest relative. Notably, a nitrogen fixation gene cluster comprising 28 genes, similar to the *Klebsiella* spp. nitrogenase gene cluster, was found in the genome of the seven pineapple strains. Based on polyphasic approaches, including ANI, dDDH, biochemical, physiological, and phylogenomic analyses, we propose the reclassification in a new species of the five pineapple strains from Hawaii A5391, A5410^T^, A5611, A6136, and A6137, together with the two pineapple strains from Malaysia CFBP 1272 and CFBP 1278, previously classified as *D. zeae*. We propose the name *Dickeya ananae* sp. nov. for this taxon, represented by the type strain A5410^T^ (= ICMP 25020^T^ = LMG 33197^T^).

## Introduction

The genus *Dickeya*, belonging to the order *Enterobacteriales* and the family *Pectobacteriaceae*, comprises an important group of pectinolytic plant pathogenic bacteria. These bacteria affect and have been isolated from a wide range of hosts, including vegetable crops, grains, and ornamental plants and also from environmental sources [1–8]. The pathogens can adapt to a wide variety of hosts and thrive in different climatic regions. These phytopathogens are notorious for causing soft rot, blackleg, and wilt in a broad array of agronomically significant crops, including potatoes, chicory, rice, cabbage, and ornamental plants, leading to substantial losses. These pectinolytic bacteria produce and secrete through type I or II secretion systems a wide array of plant cell wall-degrading enzymes (PCWDEs) [8–11] that cause maceration of the plant tissues, releasing their contents contributing to bacterial growth in the host plant [12,13]. In some pectinolytic bacteria, such as *D. dadantii* and *D. zeae*, type III secretion system has also been reported to contribute to pathogenesis [14–16]. Additionally, other virulence factors, including motility, biofilm formation, lipopolysaccharides, quorum sensing, siderophores, and the CRISPR-Cas system, contribute to the pathogenicity of these bacteria [8, 10, 14, 17–19].

The genus *Dickeya* was first established by Samson in 2005, comprising six recognized species: *D. chrysanthemi, D. dianthicola, D. dadantii, D. dieffenbachiae, D. zeae*, and *D. paradisiaca* [20]. Later, *D. dieffenbachiae* was reclassified as a subspecies of *D. dadantii* [21]. In 2014, the novel species *D. solani* was described as a predominant bacterial pathogen on potatoes in many European countries and was also isolated from hyacinths [22–24]. Recent reclassification proposed reassigning *D. paradisiaca* to the genus level and renaming it *Musicola paradisiaca* [25]. The strains gathered into the “*D. zeae* complex” exhibit high diversity, prompting further examination of this species complex [6]. In 2020, a divergent group of strains from *D. zeae* was identified as a new species, *D. oryzae* [26]. Additionally, another species emerged from the *D. zeae* complex, classified as *D. parazeae* [5]. However, this complex has not been fully resolved and is still considered a mixture of more than one species [6].

To date, the genus *Dickeya* comprises 12 species, with the type strain of each species isolated from infected plants or environmental sources. *D. dianthicola* was isolated from *Dianthus caryophyllus*; *D. dadantii* from *Pelargonium capitatum*; *D. chrysanthemi* from *Chrysanthemum morifolium*; *D. zeae* from *Zea mays*; *D. solani* from *Solanum tuberosum*; *D. fangzhongdai* from *Pyrus pyrifolia*; *D. poaceiphila* from *Saccharum officinarum*; and *D. oryzae* from rice. Three species —*D. aquatica, D. lacustris*, and *D. undicola*— were isolated from water. Additionally, the type strain of *D. parazeae*, recently separated from *D. zeae*, was also isolated from water [5]. Another proposed species, “*D. colocasiae”*, has been isolated from taro but has not yet been validly published [27].

In this study, we report the description of a new species, including the strain A5410^T^ [10], and other strains isolated from pineapples in Hawaii, causing bacterial heart rot of pineapple. This disease was first reported in Malaysia in 1957 [28] and later described in Costa Rica, Brazil, and the Philippines [29, 30]. The first outbreak in Hawaiian pineapple plantations was reported in 2003 [31], with symptoms including water-soaked lesions around the apical meristem, brown streaks on the lamina and mesophyll tissues, and characteristic blister-like lesions with light brown exudate appearing as leaves began to rot. In advanced infection stages, the pineapple heart and stem can easily separate from the plant, and young plants may be latently infected, leading to fruit collapse at maturity. The bacterium spreads through wind and water splash, with insects like pineapple fruit mites (*Steneotarsonemus ananas*) also disseminating the pathogen from infected to healthy fruits. The strains causing this disease were identified as *Erwinia chrysanthemi* (previous name of *Dickeya*) [31]. In 2008, the strain A5410 was isolated from Hawaiian pineapples and initially classified under *D. zeae* [10]. Due to the heterogeneity of *D. zeae* strains, further resolution of this species is needed [5, 6].

A recent phylogenetic study identified two major clades within the *Dickeya* genus: the first includes members of the *D. zeae* complex, and the second, termed the “main *Dickeya* clade,” comprises species like *D. dadantii, D. dianthicola* and *D. solani* [6]. Strain A5410 forms a distinct branch with dDDH similarities of 66.5%, 58.2% and 57.2% with the *type* strains of *D. oryzae*, *D. zeae*, and *D. parazeae*, respectively, supporting its unique status in a recent study of the genus *Dickeya* [6]. The genome sequence of strain A5410 was previously reported (GenBank accession number CP040816 with BioProject: PRJNA544764 and BioSample: SAMN11855518) [10] and on the basis of comprehensive whole genome comparative analysis this strain was identified as a potential candidate for a novel species. In the present research, we included six other strains isolated from pineapple and presented compelling evidence supporting the classification as a new species of seven pineapple strains isolated in Hawaii or in Malaysia. We conducted a comprehensive analysis, employing polyphasic phylogenetic analysis utilizing the *gapA* gene, 16S rRNA gene, MLSA, whole-genome sequencing, as well as phenotypic, biochemical and physiological characterization. Additionally, we used overall genome relatedness indices (OGRI) to determine genome sequence relatedness and analyzed the phenotypic characteristics of the strains. These data confirmed that the sequenced pineapple strains should be assigned to a new species of *Dickeya*. Therefore, we propose that a novel species name be established for this group of strains, *Dickeya ananae* sp. nov.

## Methods

### Bacterial strains, isolation and ecology

The strains A5391, A5410^T^, A5611, A6136, and A6137 were isolated in 2007 on Oahu (21.31 N, 158.02 W), Hawaii, from pineapple leaves (*Ananas comosus*) showing symptoms of pineapple heart rot. The bacterium was isolated by culturing on 2,3,5-triphenyltetrazolium chloride (TZC) agar medium, which contains dextrose (5 gL¹), peptone (10 gL¹), 0.001% sterilized TZC, and agar (18 gL¹) as described by Norman & Alvarez [32]. The plates were incubated at 28°C for 24 hours. A single colony was streaked onto DPA (dextrose peptone agar: DPA-dextrose 5 gL¹, peptone 10 gL¹, and agar 18 gL¹) and incubated at 28°C for 24 hours. Additionally, the colonies were streaked on crystal violet pectate (CVP) medium [33] to observe pit formation. The purified single colonies were stored at −80°C in a 30% glycerol suspension. Two additional strains isolated from pineapple in Malaysia in 1961, CFBP 1278 (NCPPB 1121, A5417) and CFBP 1272 (NCPPB 1125, A5421), were included in the study. All strains were routinely cultured on DPA media at 28°C.

### 16S rRNA and gapA gene phylogeny

The bacterial genomic DNA was isolated using the DNeasy Blood and Tissue Kit (Qiagen, Germantown, MA). For strain identification, the 16S rRNA gene was PCR amplified using the universal primers 27F and 1492R and recommended PCR conditions [34]. The PCR mix consisted of 10 µL of 2x GoTaq Green Master Mix, 1 µL of each primer (27F and 1492R; 5 µM), 1 µL of genomic DNA template, and 7 µL of nuclease-free water. Five microliters of the amplified product were enzymatically cleaned using 1 µL of Exo (exonuclease I) and 1 µL of SAP (shrimp alkaline phosphatase; GE Healthcare, Little Chalfont, UK), following the manufacturer’s instructions. The amplicons were then subjected to bi-directional DNA sequencing (Sanger method) at the GENEWIZ facility (Genewiz, La Jolla, CA). The obtained sense and anti-sense sequences were aligned and manually edited for accuracy using Geneious Prime 2023.2.1 (Biomatters, Inc., Boston, MA). The consensus sequence was compared and mapped to the whole genome of each strain and also with the NCBI GenBank nucleotide and genome databases using the BLASTn tool. The sequences of the partial 16S rRNA from A5410^T^ and other strains have been deposited in the NCBI under the accession number: [OR271567; Supplementary File 1]. The 16S rRNA phylogenetic tree was later constructed by uploading the whole genome sequences of the seven strains used in this study, along with 37 whole genomes of other *Dickeya* species, to the Type Strain Genome Server (TYGS at https://tygs.dsmz.de). The 16S rRNA phylogenetic tree was generated using the TYGS and was visualized using the web-based tool Interactive Tree Of Life (iTOL v6, https://itol.embl.de) [35].

Amplification of the partial *gapA* gene, commonly employed for strain classification within the genus *Dickeya* [36, 37], was carried out using gapA primers. The amplified product was enzymatically cleaned as previously explained. The amplicons were subjected to bi-directional DNA sequencing at the GENEWIZ facility. Obtained sense and anti-sense sequences were aligned and manually edited for accuracy using Geneious Prime. The consensus sequence was compared and mapped to the reference whole genome of each strain, as well as with the NCBI GenBank nucleotide and genome databases using the BLASTn tool. The *gapA* gene based phylogenetic tree was reconstructed using the Maximum Likelihood method and the Tamura-Nei model in MEGA 11 [38,39]. The stability of the phylogenetic tree’s topology was assessed through a bootstrap analysis with 1,000 replications.

### Genome sequencing, assembly, overall genome relatedness indices, MLSA, 16S rRNA phylogeny and phylogenomics

Purified bacterial colonies were used for genomic DNA extraction using the QIAGEN Genomic-tip 100/G (Qiagen, Valencia, CA), following the manufacturer’s instructions. The genome of the type strain A5410^T^, as reported by Boluk *et al*. [10] is available at the NCBI under the accession number CP040816. In the process of taxonomic reclassification, the genome of four additional Hawaiian strains —A5391, A5611, A6316, and A6317— as well as two strains previously reported from Malaysia, CFBP 1278 and CFBP 1272, were sequenced. The bacterial genomic DNA was extracted using the previously mentioned method, quantified with Qubit 4 using the Qubit dsDNA High Sensitivity (HS) Kit (Invitrogen, Carlsbad, CA), and analyzed on a 1.5% agarose gel. The libraries were generated without shearing to optimize sequencing read length, utilizing the Native Barcoding Kit 24 (SQK-NBD114-24) and following the protocol available on the manufacturer’s website (https://nanoporetech.com/resource-centre/protocols). A barcode-indexed library was loaded onto the R10.4.1 flow cell and sequenced using the Oxford Nanopore MinION Mk1C device (Oxford Nanopore Technologies, ONT, Oxford, UK). The obtained FAST5 sequences were used for base calling and demultiplexing using GUPPY v6.3.2 on MANA, a high-performance computing cluster at the University of Hawaii at Manoa. The *De Novo* Assemble Long Reads tool of CLC Genomics Workbench v22.0 (Qiagen) was used with default parameters to assemble the genomes.

Additionally, Illumina sequencing was performed at the DNA Technologies and Expression Analysis Core Laboratory at UC Davis Genome Center (Davis, CA). Illumina DNA libraries were prepared using 10 ng of input DNA per sample with the plexWell LP384 kit (seqWell, Beverly, MA), following the manufacturer’s instructions. This kit uses two rounds of transposase reactions followed by PCR amplification to generate a pool of dual-barcoded indexed sequencing libraries. In the first tagmentation reaction, 96 samples were individually barcoded using transposase reagents loaded with 96 different barcoded oligos, adding Illumina i7 indices and fragmenting the DNA. After a 15-minute incubation at 55°C, the reaction was halted with an SDS-containing buffer, followed by a 10-minute incubation at 68°C. The samples were then pooled in groups of 48 and purified using SPRI-bead purification with MAGwise beads. After elution in TRIS buffer, a second tagmentation was performed on the two pools, adding Illumina i5 indices to identify the sample plate. The second reaction followed similar incubation, termination, and elution steps as the first. The samples then underwent a fill-in reaction and PCR amplification, resulting in a pool of 96 sequencing libraries. The fragment length of the library pool was analyzed using a Bioanalyzer 2100 instrument (Agilent, Santa Clara, CA). The library pool was quantified by qPCR using a Kapa Library Quant RT-qPCR kit (Kapa/Roche, Basel, Switzerland) and sequenced on an Illumina NovaSeq 6000 instrument with paired-end 150 bp reads (Illumina, San Diego, CA). The Illumina paired-end short reads, along with the corrected long reads from Oxford Nanopore, were used to generate precise and comprehensive hybrid assemblies utilizing the “Polish with Reads” tool in CLC Genomics Workbench v22.0 (Qiagen) with default parameters. Contamination checks for the genomes were subsequently performed on the BV-BRC 3.26.4 web server (https://www.bv-brc.org/). Genome annotations were conducted using the Prokaryotic Genome Annotation Pipeline (PGAP v4.10) on NCBI [40] and also via the Rapid Annotation using Subsystem Technology (RAST v2.0) web server [41]. The genomes were submitted to NCBI under the reference numbers: CP040816 (strain A5410^T^), CP146223 (strain A5391), CP146227 (strain CFBP 1278), CP146228 (CFBP 1272), CP146224 (strain A6136), CP146225 (strain A6137), and CP146226 (strain A5611).

The whole genome sequences of the seven new strains were compared to that of the type strains of the 12 recognized *Dickeya* species using overall genome relatedness indices (OGRI), including ANI calculated with the OrthoANI algorithm implemented in Orthologous Average Nucleotide Identity Tool (OAT) software [42], alignment percentage (AP) using CLC Genomics Workbench 22.0.2 (Qiagen), and dDDH calculated using the Genome-to-Genome Distance Calculator version 3.0 (http://ggdc.dsmz.de/) [43], with the recommended formula 2. The calculated matrix with values for ANI and AP, as well as ANI and dDDH was generated. Additionally, OGRI was calculated between *Dickeya ananae* sp. nov., and other strains available in the NCBI database under the names *D. zeae, D. oryzae*, and *D. parazeae*.

To determine the phylogenetic position of A5410^T^, a whole genome-based taxonomic analysis was performed using the Genome BLAST Distance Phylogeny (GBDP) approach by uploading the whole genome sequences to the Type Strain Genome Server (TYGS), a free bioinformatics platform available at https://tygs.dsmz.de [43]. The distinctiveness of these strains within the genus *Dickeya* was corroborated through multi-locus sequence analysis (MLSA) [44–45], using a methodology similar to that outlined for *D. undicola* [26,46]. The sequences of ten housekeeping genes, including *dnaA, dnaX, fusA, gyrB, infB, recA, recN, rplB, atpD*, and *gapA*, were extracted from the whole genomes of these strains and other *Dickeya* species available in the NCBI database. Each sequence was aligned using Geneious Prime, and a phylogenetic tree was constructed using MEGA 11. Evolutionary history was inferred using the Maximum Likelihood method and the Tamura-Nei model, with 1,000 bootstrap replicates performed.

### Pan and core genome analyses

Pan- and core-genome analyses were performed using the whole genome sequences of the newly described species and type strains of closely related species within the genus *Dickeya*. Annotated files (GFF3) were generated using Prokka v1.14.6 [47] and used for pan-genome analysis with the Roary v3.13.0 pipeline [48], employing a minimum BLASTp identity of 90%. Core genomes were aligned using PRANK, a probabilistic multiple alignment program [48,49]. Core and unique gene counts for the type strain of each *Dickeya* species were determined from the Roary output (with a BLASTp identity threshold of 80%), and these data were used to create flower plots using R scripts in RStudio. A core gene phylogenetic tree was constructed using the ML tree inference tool RAxML-NG v0.8.0 (Randomized Axelerated Maximum Likelihood - Next Generation) [50]. The DNA substitution model, General Time Reversible (GTR) + GAMMA (G), was applied separately to the core genomes of the *Dickeya* species, with 1,000 bootstraps. The resulting core genome phylogenetic tree was visualized using iTOL. The Roary matrix, which delineates the presence and absence of core and accessory genes, was combined with the core genome maximum likelihood tree, and the results were presented using roary_plots.py [48].

### Phenotypic characteristics of the pineapple strains

The bacteria were tested for swimming and swarming motilities using a semisolid medium [51]. The swimming assay medium contained 10 g/L tryptone, 5 g/L NaCl, and 3 g/L agar, while the swarming medium had 10 g/L tryptone, 5 g/L NaCl, and 4 g/L agar. Five microliters of overnight culture with an OD at 600 nm of 1.0 were spot inoculated at the center of each medium plate, and the diameter of bacterial growth was recorded. Each assay was performed in triplicate. The GEN III MicroPlate (BIOLOG) was used to assess carbon source utilization and chemical sensitivity of strains A5391, A5410^T^, A5611, A6316, A6317, CFBP 1278 (A5417), and CFBP 1272 (A5421), following the manufacturer’s instructions. The bacterial suspension was prepared in inoculating fluid (IF) by suspending pure bacterial colonies from overnight cultures grown on BUG (Biolog Universal Growth) media, adjusted to 96% transmittance using a SPECTRONIC 20D+ spectrophotometer (Thermo Fisher Scientific, MA, USA). A homogeneous suspension was prepared, and 100 µl of the suspension was dispensed into each well of the Biolog GEN III MicroPlate. The plates were incubated at 28 °C for 24 hours, read in the Biolog Microstation ELx808BLG Reader (Biotek Instruments), and the results were recorded as utilization or no utilization for carbon sources, and sensitivity or resistance for chemicals.

Biochemical test kits API 20E, APIZYM, and API 50CH from bioMérieux (Marcy-l’Étoile, France) were used to investigate carbohydrate fermentation, biochemical reactions, and enzymatic activities. The bacteria were cultured on NAG media, and 24-hour plates were used for the tests. For the API 20E strips, which consist of 21 miniaturized biochemical tests, the bacterial inoculum was prepared by suspending a single isolated colony in API suspension medium (0.85% NaCl). The inoculum was then used immediately to fill the test tubes and cupules, following the manufacturer’s instructions. The test was incubated at 28±2°C, and results were recorded after 24 hours. For the API ZYM enzymatic assay, which identifies 19 enzymatic reactions semi-quantitatively, the bacterial inoculum was prepared in API suspension medium with turbidity adjusted to 5 McFarland. Then, 65 µl of the suspension was added to each cupule and incubated at 28±2°C for 4 hours and 30 minutes. The test was developed by adding one drop each of ZYM A and ZYM B reagents and observed after 5 minutes for color changes to determine enzyme activity. API 50CH strips, designed to test the utilization of 49 different carbohydrate sources, were used following the *Enterobacteriaceae* instructions [52]. Briefly, colonies from 18-24 hour-old NAG plates were suspended in sterile distilled water, adjusted to 4 McFarland, mixed with CHB/E medium, and used to inoculate the strips. Strips were incubated at 28±2°C and results were observed at 24 and 48 hours. Cytochrome oxidase activity was tested using OxiStrips^TM^ (Hardy diagnostics, Santa Maria, CA), with a color change to dark blue within 90 seconds indicating a positive result. Catalase activity was confirmed by the addition of 3% H_2_O_2_, with bubble formation indicating a positive result.

Extracellular enzyme production, including pectate lyase (pel), cellulase (cel), and protease (prt), was assessed using a semi-quantitative method [53]. Assays involved applying 50 µl of a bacterial suspension with OD600 ∼1.2 to wells in media plates and incubating at 28°C. The pel assay was developed with 4N HCl after 10 hours, the cel activity with 2% Congo red and 5 M NaCl, and the prt activity observed as a halo after 24 hours of incubation.

The maceration ability of these strains was evaluated through virulence assays on potato, taro, and onion. The surface of each tuber or bulb was sterilized with 0.06% sodium hypochlorite for 10 minutes, followed by three 3-minute washes in sterile distilled water. The tubers and bulbs were air-dried in a BSC II, sliced with a sterile knife, and placed in sterile Petri plates. A 10 µl bacterial suspension of 10^8^ CFU/ml was applied to the center of each slice, and plates were incubated at room temperature (26°C). Results were recorded after 24 hours, with two replicates per strain.

### Transmission Electron Microscopy (TEM)

To examine bacterial morphology, transmission electron microscopy (TEM) was conducted. Samples were prepared following the protocol from the Biological Electron Microscope Facility, University of Hawaii at Manoa. Bacteria were cultured overnight in Tryptic Soy broth at 28°C with shaking at 120 rpm. For TEM, bacteria were mounted onto glow-discharged 200 mesh carbon-coated Formvar-coated copper grids, negatively stained with 1% uranyl acetate, and rinsed thrice with sterile distilled water. Stained grids were viewed on a Hitachi HT7700 TEM at 100 kV, and high-resolution images were captured using an AMT BioSprint16M-ActiveVu: 16 megapixel cooled 4895 x 3920-pixel CCD camera. Cells were measured in length and diameter with three replicates.

### Antibiotic sensitivity assay

An antibiotic disc diffusion assay was performed. Briefly, a single colony from a 12-hour grown culture plate was inoculated into 10 ml of nutrient broth supplemented with 0.2% glucose and incubated at 28°C with shaking at 150 rpm for 8 hours. The bacterial inoculum was adjusted to an OD_600_ of 1.0, and 100 µl was evenly spread onto NAG plates. The plates were allowed to dry in the biosafety cabinet. The antibiotic discs were prepared by absorbing discs with the following 13 antibiotics: spectinomycin (100 mg/ml), gentamicin (50 mg/ml), kanamycin (50 mg/ml), tetracycline (40 mg/ml), penicillin (50 mg/ml), cefalexin (50 mg/ml), chloramphenicol (50 mg/ml), trimethoprim (25 mg/ml), bacitracin (50 mg/ml), hygromycin B (50 mg/ml), vancomycin (50 mg/ml), cefotaxime (50 mg/ml), and carbenicillin (50 mg/ml). Once the discs were dried, one disc was placed at the center of each agar plate by gently pressing down with a sterile tweezer; 2 plates were prepared for each antibiotic tested. For the control, a disc impregnated with sterile distilled water was placed in the center of the culture plate. The plates were inverted and incubated for 24 hours at 28°C. The area of inhibited bacterial growth was measured and recorded after 24 hours.

## Results and Discussion

### Genomic features and genomic relatedness

The genome of strain A5410^T^ was sequenced by Boluk *et al.* [10] and used for comparative genomic analyses. This study provided evidence that the taxonomic position of this pineapple strain should be revised, as also pointed out by Hugouvieux-Cotte-Pattat *et al.* [6]. In the current study, we included the bacterial strains CFBP 1278 and CFBP 1272, isolated from pineapple in Malaysia, along with strains A5391, A6136, A6137, and A5611, which were isolated in 2006 during a bacterial heart rot outbreak in Hawaiian pineapple. The strains isolated from pineapple in Hawaii were collected from the base of leaves from different pineapple samples. The six pineapple strains were sequenced using Illumina NovaSeq and MinION sequencing platforms. The hybrid assembly approach employed yielded high-quality genomes with coverage ranging from 364x to 545x. Genomic features are presented in Table 1. A CheckM-based assessment demonstrated genome completeness within the range of 99.9-100%, with contamination rates ranging from 2.2% to 4%. The genome sizes ranged 4,779,825 bp to 4,837,911 bp, with no plasmids detected. The G+C content ranged from 53.48% to 53.56 mol%. Results from the Bacterial and Viral Bioinformatics Resource Center (BV-BRC version 3.34.11) indicated that the A5410^T^ genome encodes a total of 4,555 protein-coding sequences, 22 rRNAs, 75 tRNAs, 3 CRISPR arrays, and 884 hypothetical proteins. The six other genomes of pineapple strains showed similar values (Table 1).

**Table 1:** Genome features and Statistics for the six strains of *Dickeya ananae* sp. nov. other strains sequenced in this study. The genomic features as annotated were assessed by The Bacterial and Viral Bioinformatics Resource Center (BV-BRC version 3.34.11).

|  | Strains |  |  |  |  |  |  |
| --- | --- | --- | --- | --- | --- | --- | --- |
| Genome Statistics | A5410<sup>T</sup>* | A5391 | CFBP 1278 (A5417) | CFBP 1272 (A5421) | A6136 | A6137 | A5611 |
| GenBank accession number | CP040816.1 | CP146223 | CP146227 | CP146228 | CP146224 | CP146225 | CP146226 |
| Biosample | SAMN11855518 | SAMN39956632 | SAMN39957828 | SAMN39957829 | SAMN39956949 | SAMN39956997 | SAMN39956998 |
| Bioproject | PRJNA544764 | PRJNA1076913 | PRJNA1076913 | PRJNA1076913 | PRJNA1076913 | PRJNA1076913 | PRJNA1076913 |
| Assembly level | complete | complete | complete | complete | complete | complete | complete |
| Genome Sequencing Platform | PacBio RSII | Hybrid (MinIon+illumina Novaseq) | Hybrid (MinIon+illumina Novaseq) | Hybrid (MinIon+illumina Novaseq) | Hybrid (MinIon+illumina Novaseq) | Hybrid (MinIon+illumina Novaseq) | Hybrid (MinIon+illumina Novaseq) |
| Genome coverage (x) | 558.0x | 364.20x | 476.99x | 336.798x | 469.897x | 459.07x | 545.335x |
| Contigs | 1 | 1 | 1 | 1 | 1 | 1 | 1 |
| Genome Length (bp) | 4779199 | 4834333 | 4805951 | 4797088 | 4779825 | 4779844 | 4837911 |
| GC Content (mol%) | 53.481453 | 53.492363 | 53.52932 | 53.5207 | 53.48164 | 53.48093 | 53.56957 |
| Contig L50 | 1 | 1 | 1 | 1 | 1 | 1 | 1 |
| Contig N50 length (bp) | 4779199 | 4834333 | 4805951 | 4797088 | 4779825 | 4779844 | 4837911 |
| **Annotation Statistics** |  |  |  |  |  |  |  |
| No. of transfer RNAs (tRNAs) | 75 | 75 | 76 | 76 | 75 | 75 | 75 |
| No. of ribosomal RNAs (rRNAs) | 22 | 22 | 22 | 22 | 22 | 22 | 22 |
| CDS | 4555 | 4728 | 4658 | 4629 | 4692 | 4708 | 4756 |
| CDS Ratio | 0.9530886 | 0.97800463 | 0.969215 | 0.96496 | 0.981626 | 0.984969 | 0.983069 |
| Hypothetical CDS | 884 | 911 | 876 | 877 | 898 | 917 | 907 |
| Hypothetical CDS Ratio | 0.29330406 | 0.29483926 | 0.287033 | 0.289263 | 0.290494 | 0.293968 | 0.291421 |
| PLFAM CDS | 4096 | 4219 | 4194 | 4164 | 4214 | 4225 | 4229 |
| PLFAM CDS Ratio | 0.8992316 | 0.89234346 | 0.900386 | 0.899546 | 0.898124 | 0.897409 | 0.889193 |
| CRISPR_array | 3 | 3 | 2 | 2 | 3 | 3 | 3 |
| **Genome Quality** |  |  |  |  |  |  |  |
| Coarse Consistency | 99.6 | 99.5 | 99.3 | 99.6 | 99.4 | 99.6 | 99.5 |
| Fine Consistency | 99.1 | 96.6 | 97 | 97.1 | 96.8 | 96.8 | 96.4 |
| CheckM Completeness | 100 | 100 | 100 | 99.9 | 99.9 | 100 | 100 |
| CheckM Contamination | 0.3 | 3.1 | 4 | 2.2 | 2.5 | 4 | 3.6 |
| Genome Quality | Good | Good | Good | Good | Good | Good | Good |

\*Previously sequenced and reported in Boluk et al., 2021

Annotation and comparison of the genome sequences using the RASTtk Model SEED database revealed that 28-29% of the predicted coding sequences were assigned to subsystems. The number of subsystems predicted by the RAST annotation server ranged from 342 to 345 (Supplementary Table 1). The subsystems most prominently represented among the seven genomes included “amino acids and derivatives”, along with “carbohydrates”. Additional common subsystems were associated with “protein metabolism”, “cofactors, vitamins, prosthetic groups, pigments”, and “respiration”.

A comprehensive comparative genomics analysis conducted by Boluk *et al.* [10] identified a unique nitrogen fixation cluster in the A5410^T^ genome, which was absent in the strains of *D. oryzae*. The six other genomes sequenced in this study contain a similar cluster, including those from Malaysia. The *nif* gene cluster was absent in all type strains of the genus *Dickeya*. It was also absent in strains within the *D. zeae* species complex but was present in strains CE1 (isolated from *Canna*, China), JZL7 (isolated from *Clivia*, China), and BRIP 64263 (isolated from pineapple, Australia). (Figure 1). The *nif* gene cluster of the A5410^T^ genome consisted of a total of 20 genes spanning a 25,607 bp region. The arrangement of this gene cluster is similar to that of the nitrogen fixation *nif* cluster of *Klebsiella variicola* and *K. pneumoniae*, comprising the three structural genes: *nifH* (nitrogenase reductase), *nifD* (nitrogenase molybdenum-iron protein α subunit), and *nifK* (nitrogenase molybdenum-iron protein β subunit). It also includes *nifF* and *nifJ* (electron donation to nitrogenase); *nifE, nifN, nifX, nifU, nifS,* and *nifV* (Fe-Mo cofactor biosynthesis); *nifM* (nitrogenase maturation); *nifA* and *nifL* (positive and negative regulators of *nif* gene expression); *nifW* (protection, maturation, and activation of the Fe-Mo protein); and *nifT* of unknown function [54–58]. An additional gene encoding a hypothetical protein is present between *nifJ* and *nifH* in the *nif* gene cluster of the seven pineapple strains (Figure 1).

**Figure 1.**
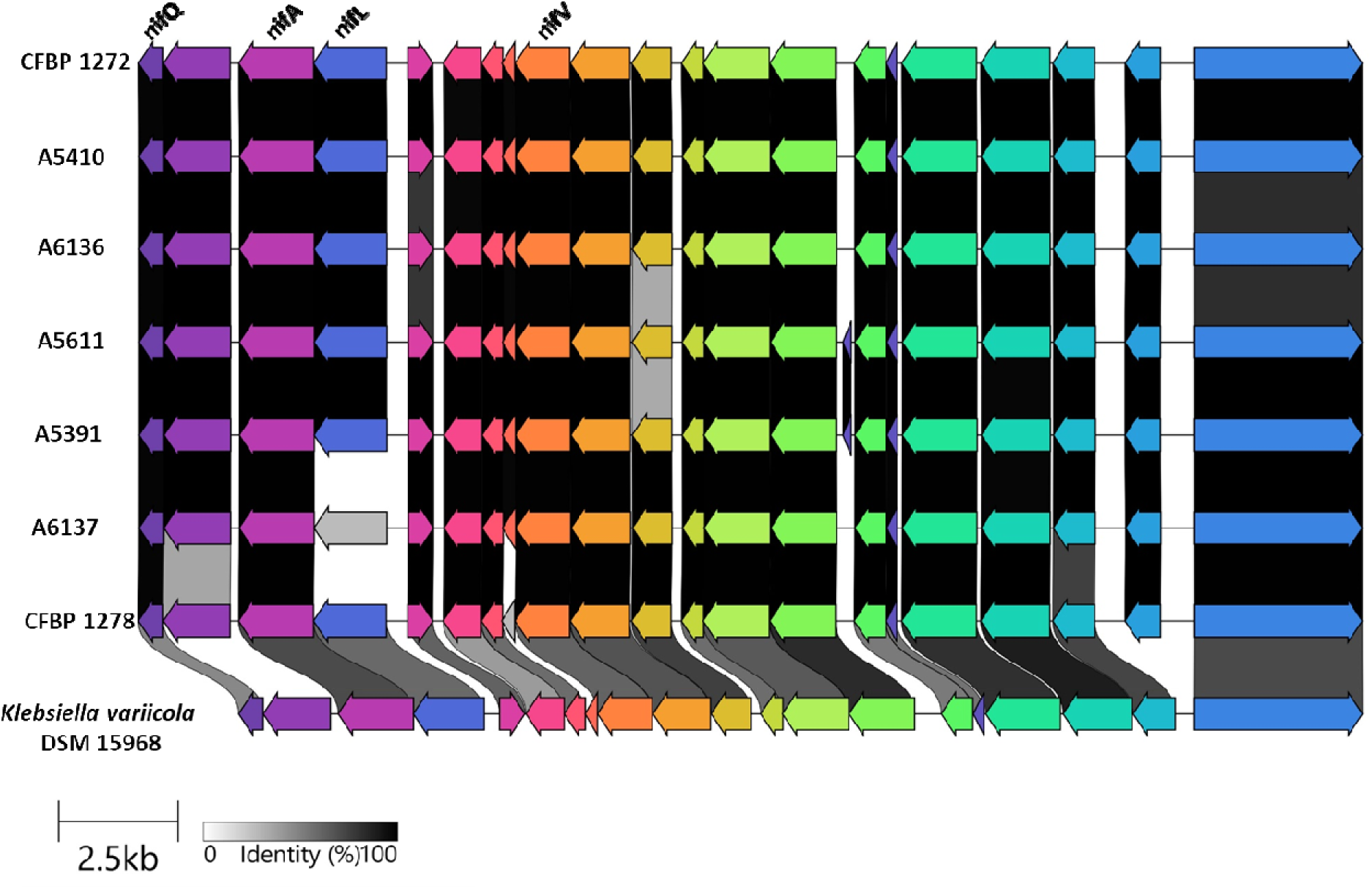
Comparison of the genetic organization of the nitrogen fixation (*nif*) gene cluster among *Dickeya ananae* sp. nov. strains and *Klebsiella variicola* DSM 15968. The arrow positions indicating forward or reverse gene orientation. Arrow colors represent the specific gene composition within the *nif* gene cluster. Gene names are displayed at the top and bottom of the linear graph. Pairwise alignment of the sequences was conducted using the BLAST algorithm with cut-off values of 70%-100%, with regions of higher nucleotide identity shaded in grey and black. Expanded legend entries acronyms are provided: *nif Q* (Fe-Mo cofactor synthesis), *nif B* (Fe-Mo cofactor synthesis), *nifA* (transcriptional activator), *nifL* (transcriptional regulator), *nifF* (electron donation to nitrogenase), *nifM* (nitrogenase maturation), *nifZ* (activation and maturation), *nifW* (protection of the Fe-Mo protein; maturation and activation, *nifV* (Fe-Mo cofactor synthesis), *nifS* (cysteine desulfurase; maturation and activation), *nifU* (Fe-Mo cofactor biosynthesis), *nifX* (Fe-Mo cofactor biosynthesis), *nifN* (Fe-Mo cofactor biosynthesis), *nifE* (Fe-Mo cofactor biosynthesis), *nifY* (Fe-Mo cofactor synthesis), nifT (unknown function), nifK (nitrogenase molybdenum-iron protein β subunit), nifD (nitrogenase molybdenum-iron protein α subunit), *nifH* (nitrogenase reductase), HP (hypothetical protein), *nifJ* (Fe electron transport).

Plant cell wall-degrading enzymes, including pectinases, cellulases, and proteases, are among the most important virulence factors of soft rot *Pectobacteriaceae*. The genome of A5410^T^ was previously analyzed for virulence-related genes by Boluk *et al.* [10], and in this study, we also evaluated the presence of these genes in the six other pineapple strains. Genes encoding pectate lyase (*pelADE, pelBCZ, pelI, pelL, pelN, pelX*, and *pelW)*; other pectinases (*pemA, paeY, faeD, pehK, pehX, rhiE*); a cellulases (*celZ*); proteases (*prtABCDEFG*); a xylanase (*xynA*); and a phospholipase (*plcA*) were present in the seven genomes. The genes *pnlH*, *pehN*, and *pemB, and avrM* encoding a pectin lyase, a polygalacturonase, a pectin methylesterase, and an avirulence protein respectively, are absent in these genomes, as well as in *D. zeae, D. parazeae* and *D. oryzae* [6]. All the genomes of the proposed *D. ananae* sp. nov. members possess the *avrL* avirulence-associated gene, unlike the *D. oryzae* strains ZYY5^T^, DZ2Q, and EC1 [5]. The *pnlG,* which encodes *a* pectin lyase, is absent in *D. ananae* members as well as *D. parazeae,* but is present in *D. oryzae* and *D. zeae,* as noted by Hugouvieux-Cotte-Pattat *et al*. [5].

Furthermore, the genomes of the proposed *D. ananae* sp. nov. members also contained *cyt* genes encoding Type-1Ba cytolytic delta-endotoxin and Type-2Aa cytolytic delta-endotoxin, which were absent in *D. oryzae* ZYY5^T^, DZ2Q, and EC1. The *cyt* genes were reported to be present in some, but not all, strains of *D. oryzae* [5].

### Genomic Relatedness

Whole genome alignment and 16S rRNA sequence analysis performed using the TYGS genome server identified A5410^T^ and the six other pineapple strains as a novel species, all belonging to the same group. The taxonomic position of the pineapple strains was first clarified by comparing the pairwise ANI and dDDH values between the A5410^T^ genome and the type strains of each *Dickeya* species (Figure 2). These ANI and dDDH values ranged from 79.6-96% and 23.1-66.8%, respectively, falling below or equal to the thresholds of 95–96% (ANI) and 70% (dDDH) for differentiating bacterial species, confirming the taxonomic novelty of strain A5410^T^ [59–62]. The ANI values between the A5410^T^ genome and the other six genomes sequenced in this study (strains A6136, A6137, A5611, A5391, CFBP 1272, and CFBP 1278) ranged from 99.01-99.99% (Supplementary Table 2). Additionally, the dDDH values between these seven genomes ranged from 91.8-99.9%, strongly supporting that all seven strains belong to the same species. Alignment coverage, represented by AP values between genomes, is one of the criteria evaluated in the description of new species by Parsenen *et al.* [63]. The AP values between the genomes of A5410^T^ and the six other strains ranged from 97.56-99.42%, strongly supporting their genomic similarity and classification within the same group. In comparison, AP values between the A5410^T^ genome and those of type strains of each *Dickeya* species range from 26.7 (with *D. lacustris*) to 87.2 (with *D. zeae*) (Figure 3).

**Figure 2.**
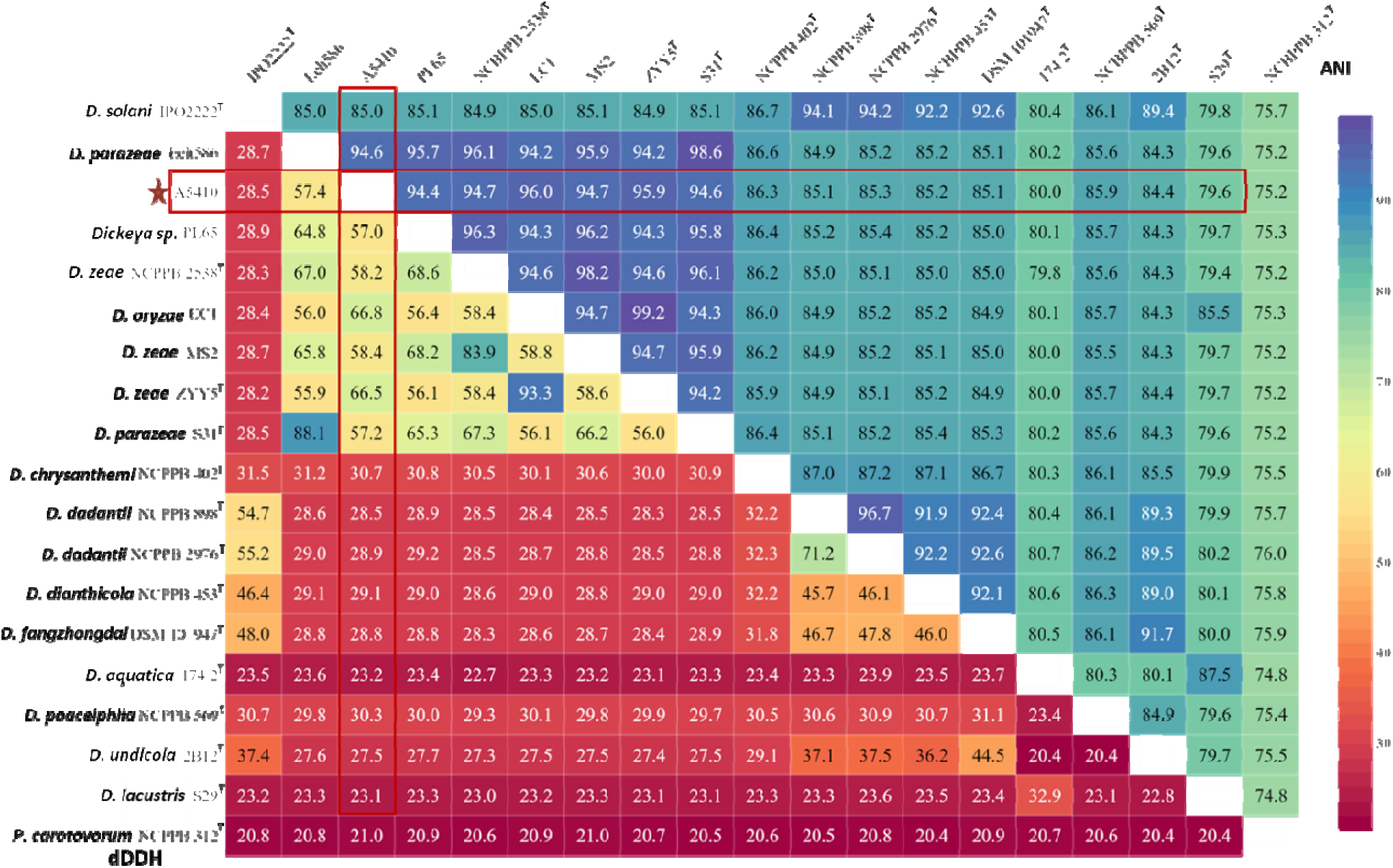
Heatmap displaying combined ANI and dDDH values. ANI values (above diagonal) were calculated using the OrthoANI algorithm in OAT software; dDDH values (below diagonal) were determined using the Genome-to-Genome Distance Calculator (GGDC) version 3.0 (https://ggdc.dsmz.de/). The analysis includes the type strains of each *Dickeya* species and *D. ananae* sp. nov. strain A5410^T^.

**Figure 3.**
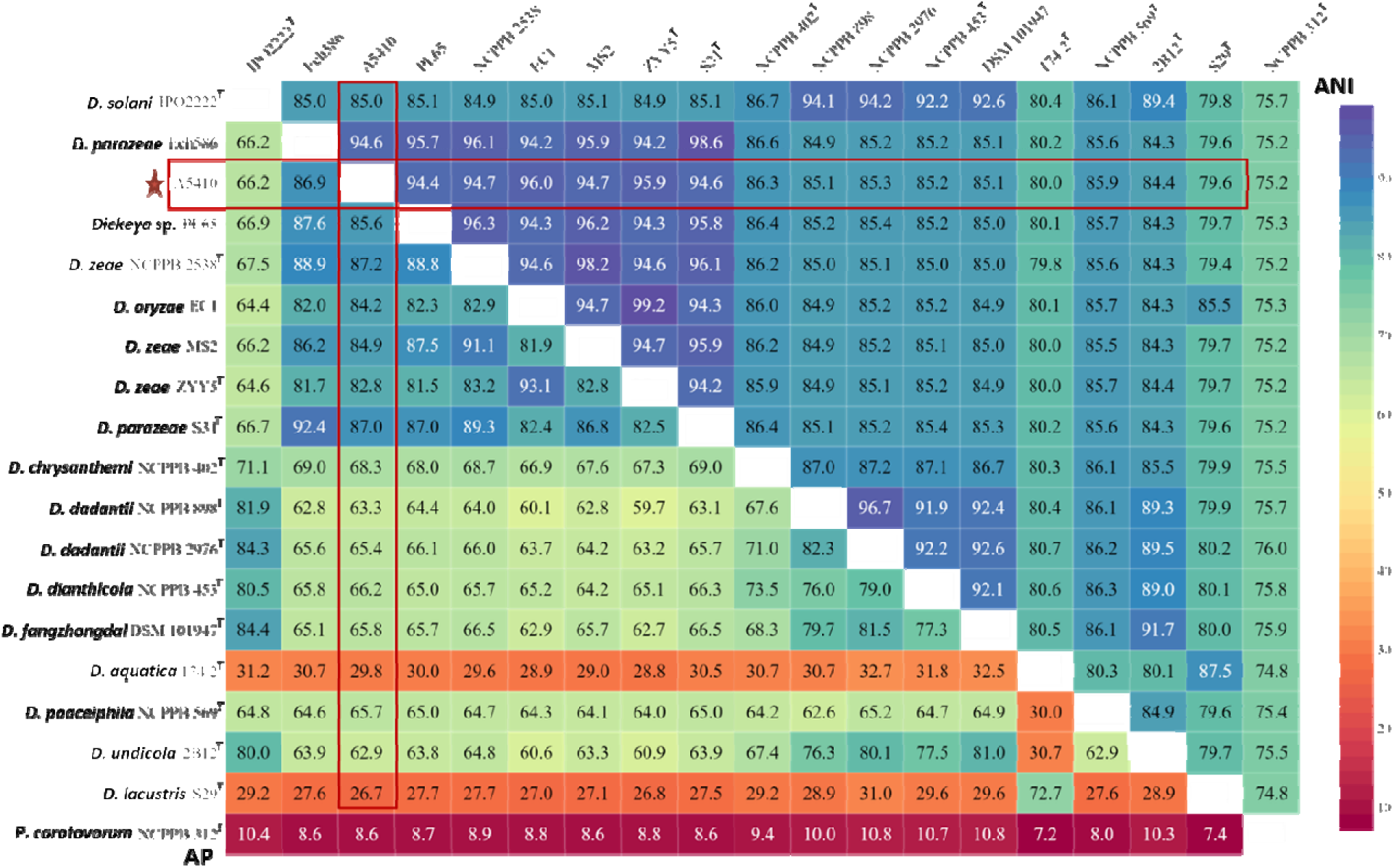
Heatmap showing the combined ANI and AP values. ANI values (above diagonal) and AP values (below diagonal) were calculated using CLC Genomics workbench 22.0.2 for all type strains of the genus *Dickeya* and *D. ananae* sp. nov. A5410^T^.

Given the reported diversity among members of the *D. zeae* complex [5,6], the genomes of the seven pineapple strains were further evaluated for ANI and dDDH values in comparison to strains of closely related groups within the *D. zeae* complex, including the species *D. parazeae, D. zeae,* and *D. oryzae* (Supplementary Table 2). These genomes share ANI and dDDH values of 94.47-94.62% and 57-57.5%, respectively, with the genomes of the three *D. parazeae* strains S31^T^, Ech 586 and A586-S18-A17. Comparison with the genomes of ten *D. zeae* strains (NCPPB2538^T^, JZL7, PL65, CE1, MS1, MS2014, MS2018, MS2, MK19, NCPPB3532) showed ANI and dDDH values ranging from 94.34-94.79% and 56.7-59%, respectively (Supplementary Table 2). The ANI and dDDH values between the genomes of the seven pineapple strains with the genome of the *D. oryzae* type strain ZYY5^T^ range from 95.93-96% and 66.4-66.5%, respectively. Additional strains reported to be members of, or close to, *D. oryzae* were also evaluated for ANI and dDDH values with the seven pineapple strains. As described by Hugouvieux-Cotte-Pattat *et al*. [5], the strains ZYY5^T^, DZ2Q, ZYU1202, and EC1 were reported to be more closely related to each other than to EC2 (all five strains were isolated from rice). The calculated ANI and dDDH values were 95.97-96% and 66.4-66.9%, respectively, with the genomes of EC1, ZYU1202, or DZ2Q, and reached 96.2-96.4% and of 69.1-69.5%, respectively, with the EC2 genome (Supplementary Table 2). The ANI values were at the boundary for species delineation, ranging from 96.1-96.6% with other strains of *D. oryzae*, such as S20, CSLRW192, and FGV03 (all isolated from water), and NCPPB 3531 (isolated from potato). dDDH values range from 68.2-69.5% with strains S20, CSLRW192, and FGV03, and 66.9-70.3% with NCPPB 3531. These values, at the limit of those recommended for the delimitation of species, show the difficulty of using only ANI and dDDH data for the description of species. The ANI values are in the same range as those reported for the recent description of the species *P. parvum* and *P. quasiaquaticum* [63,64]. However, analysis of the alignment coverage with the genomes of *D. oryzae* strains S20, CSLRW192, FGV03, and NCPPB 3531 showed AP values (85.91-89.56%) lower than those observed with other *Dickeya* species, such as *D. zeae* and *D. parazeae* (87-87.2 %) (Supplementary Table 2, Figure 3). In comparison, the genomes of the seven pineapple strains share a high alignment coverage percentage (97.03-99.42%), indicating close relatedness within them. Therefore, the analysis of ANI, dDDH, and AP values is consistent with the proposition of a novel species within the *D. zeae* complex, borderline with some *D. oryzae* strains.

### *gapA*, 16SrRNA, MLSA, whole genome and pan-genome phylogeny

The gene *gapA* (encoding glyceraldehyde-3-phosphate dehydrogenase) has been routinely used for the identification and accurate phylogenetic positioning of strains within the genus *Dickeya* [36]. Amplification of the *gapA* gene sequence confirmed that the seven pineapple strains belong to the genus *Dickeya*. Since the strains were closely grouped with members of the *D. zeae* complex, *gapA* sequences of the type strains of *D. zeae, D. parazeae, D. oryzae,* along with other strains available in the NCBI database, were extracted (Supplementary Table 3), aligned, and trimmed to 702 bp of the *gapA* gene region using Geneious Prime. This alignment was used to build a phylogenetic tree using the maximum likelihood method based on the Tamura-Nei model to infer the phylogenetic position. The *gapA* phylogenetic tree illustrates that the seven pineapple strains A5410^T^, A5391, CFBP 1278 (A5417), CFBP 1272 (A5421), A6136, A6137, and A5611 cluster together to form a distinct monophyletic clade, strongly supported by a high bootstrap percentage (Figure 4). The strains CFBP 1278 and CFBP 1272 (previously submitted as *D. zeae*), isolated from pineapple in Malaysia, also grouped within this monophyletic clade, consistent with previous MLSA studies by Marrero *et al*. [65], indicating that the strains isolated from pineapple may constitute a novel genomic species. The corresponding clade is positioned between *D. parazeae* and *D. oryzae*, closely related to *D. oryzae* but clearly distinct from it and other members of the genus [5], supporting the delineation of a new species.

**Figure 4.**
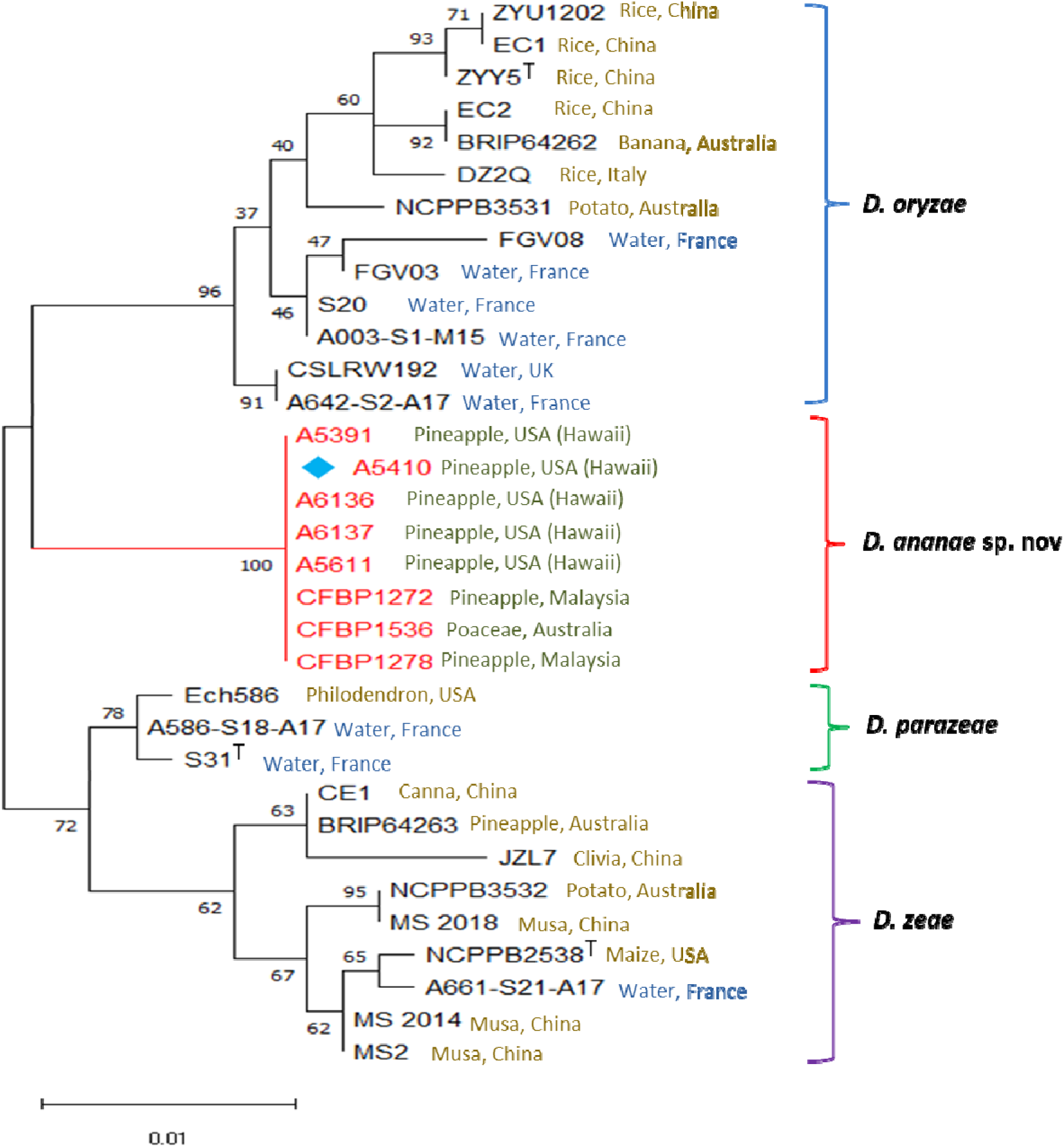
The *gapA* based phylogenetic tree of the *Dickeya* species. This tree includes different strains of each species included in the *D. zeae* complex. Evolutionary history was inferred by using the maximum likelihood method and Tamura-Nei model. The tree with the highest log likelihood (−1409.19) is shown. The percentage of trees in which the associated taxa clustered together is shown next to the branches. Initial tree(s) for the heuristic search were obtained automatically by applying Neighbor-Join and BioNJ algorithms to a matrix of pairwise distance estimated using the Tamura-Nei model, and then selecting the topology with superior log likelihood value. The tree is drawn to scale, with branch lengths measured in number of substitutions per site. This analysis involved 33 nucleotide sequences. There was a total of 702 positions in the final dataset. Evolutionary analyses were conducted in MEGA11.

The BLAST search in NCBI nearly full length of 16S rRNA gene sequence of strain A5410^T^ showed the highest sequence similarity of 98.57% with *D. oryzae* ZYY5^T^ followed by 97.80% with S31^T^ *D. parazeae* and 97.05% with *D. zeae* NCPPB 2538^T^. The partial 16S rRNA sequence of the proposed type strain A5410^T^ was deposited in the GenBank nucleotide database with the accession number OR271567, and the partial sequences of the 16S rRNA amplified using the 27F and 1492R primers [34] are provided in Supplementary File 1. The partial amplified 16S rRNA gene sequences of the *D. ananae* sp. nov. strains were further mapped to the whole genome of each sequenced strain, resulting in a 100% match. The 16S rRNA phylogenetic tree, encompassing 50 strains (including type strains within the genus *Dickeya,* other *Dickeya* sp. complex strains, strains listed as *Dickeya* sp., and *P. carotovorum* as an outgroup; Supplementary Table 3), was inferred through the Type (Strain) Genome Server (TYGS), a high-throughput platform for accurate genome-based taxonomy (https://tygs.dsmz.de). Utilizing the GBDP method and full-length 16S rRNA gene sequences without ambiguities, the phylogeny separated the *D. zeae* species complex into two clades: one clade includes the strains of the *D. oryzae, D. parazeae*, and *D. zeae* species, and another clade includes the seven *D. ananae* sp. nov. strains which clustered together as a monophyletic clade. This 16S rRNA clustering also supports that the novel species is distinct from the other members of the *D. zeae* complex (Supplementary Figure 1).

To further refine the taxonomic position of A5410^T^, MLSA based on ten housekeeping genes was conducted, and the phylogenetic positions of the strains were inferred using the Maximum Likelihood method and the Tamura-Nei model with 1,000 bootstraps. The nucleotide sequences of the ten housekeeping genes were retrieved from the NCBI Genome database, encompassing 50 strains, including each *Dickeya* species and *P. carotovorum* (used as an outgroup). Each gene sequence was trimmed and assembled to generate a consensus sequence. The ten housekeeping genes comprised *atpD* (1,369 bp), *dnaA* (1,325 bp), *dnaX* (2,058 bp), *fusA* (2,110 bp), *gapA* (1,073 bp), *gyrB* (2,490 bp), *infB* (2,722 bp), *recA* (1,083 bp), *recN* (1,655 bp), and *rplB* (807 bp), resulting in a total concatenated sequence length of 16,692 bp. This analysis clearly separated *D. ananae* sp. nov. strains into a distinct clade with a high bootstrap percentage (Figure 5). The topology of the ML tree clearly showed that the *D. zeae* species complex is divided into two major clades: the first clade includes members from *D. oryzae* and *D. ananae* sp. nov., and the second clade comprises members of *D. zeae* and *D. parazeae*. The stability of the phylogenetic tree topology was well supported by high bootstrap values, further supporting the clustering of *D. ananae* sp. nov. members in a monophyletic clade.

**Figure 5.**
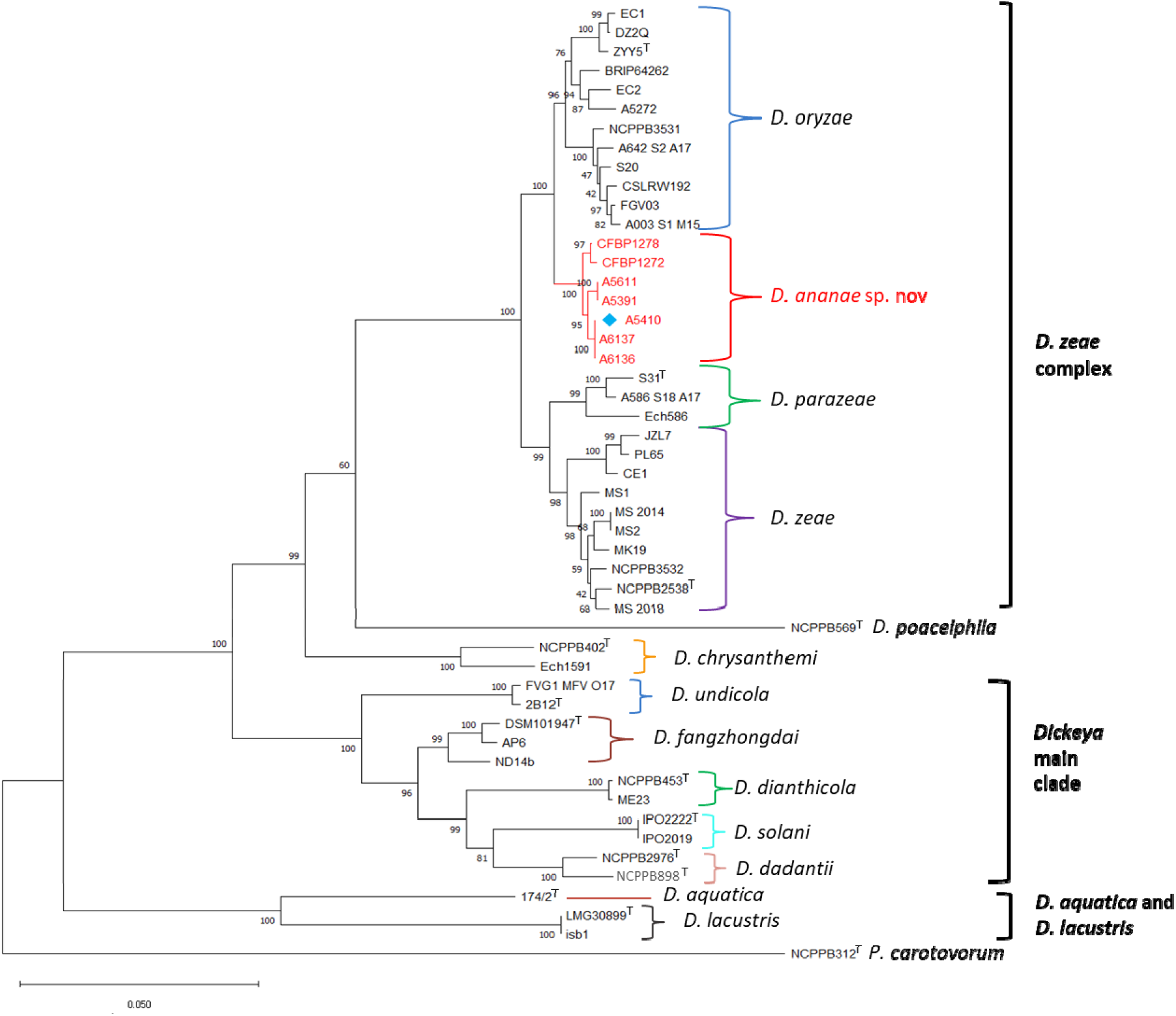
A maximum likelihood phylogenetic tree based on the concatenated sequences of 10 housekeeping genes, *dnaA*, *dnaX*, *fusA*, *gyrB*, *infB*, *recA*, *recN*, *rplB*, *atpD* and *gapA* retrieved from the whole genome sequences of 50 strains including the type and other strains of the other species within genus *Dickeya*. The multiple alignment of each gene sequence and concatenation was performed using Geneious Prime 2021.2.2. The phylogenetic tree was reconstructed using concatenated alignments in MEGA 11 with 1,000 bootstrap replicates. Evolutionary distance were calculated using the Tamura-Nei model. Bootstrap percentages are indicated at the branch points (only values above 60% are shown). The genome of *Pectobacterium carotovorum* wa used as an outgroup. The tree scale bar indicates the number of nucleotide substitutions per sequence position. The percentage of trees in which the associated taxa clustered together i shown next to the branches. Initial tree(s) for the heuristic search were obtained automatically by applying Neighbor-Join and BioNJ algorithms to a matrix of pairwise distances estimated using the Tamura-Nei model, and then selecting the topology with superior log likelihood value. The tree is drawn to scale, with branch lengths measured in the number of substitutions per site. This analysis involved 50 nucleotide sequences. There was a total of 16,692 positions in the final dataset.

The whole-genome phylogenetic tree was created using the TYGS webserver based on the Genome Blast Distance Phylogeny (GBDP) distance formula *d_5_* approach, with *P. carotovorum* included as an outgroup (Figure 6). This analysis clearly separated the strains of the *D. zeae* complex into two main clades: Clade I, which branched into two subclades corresponding to *D. oryzae* strains and *D. ananae* sp. nov. strains, respectively, and Clade II, which branched into three subclades corresponding to *D. zeae, D. parazeae,* and potential other species, with high bootstrap values supporting the tree topology. Thus, on the whole-genome phylogenetic tree, strains of *D. ananae* sp. nov. clearly group separately as a distinct monophyletic clade (Figure 6). The strains of *D. ananae* sp. nov. form a distinct and separate clade, closely related to *D. oryzae* and its members, which are divided into two clades in all phylogenetic analyses performed in this study, consistent with the description of the *D. zeae* complex by Hugouvieux-Cotte-Pattat *et al*. [5].

**Figure 6.**
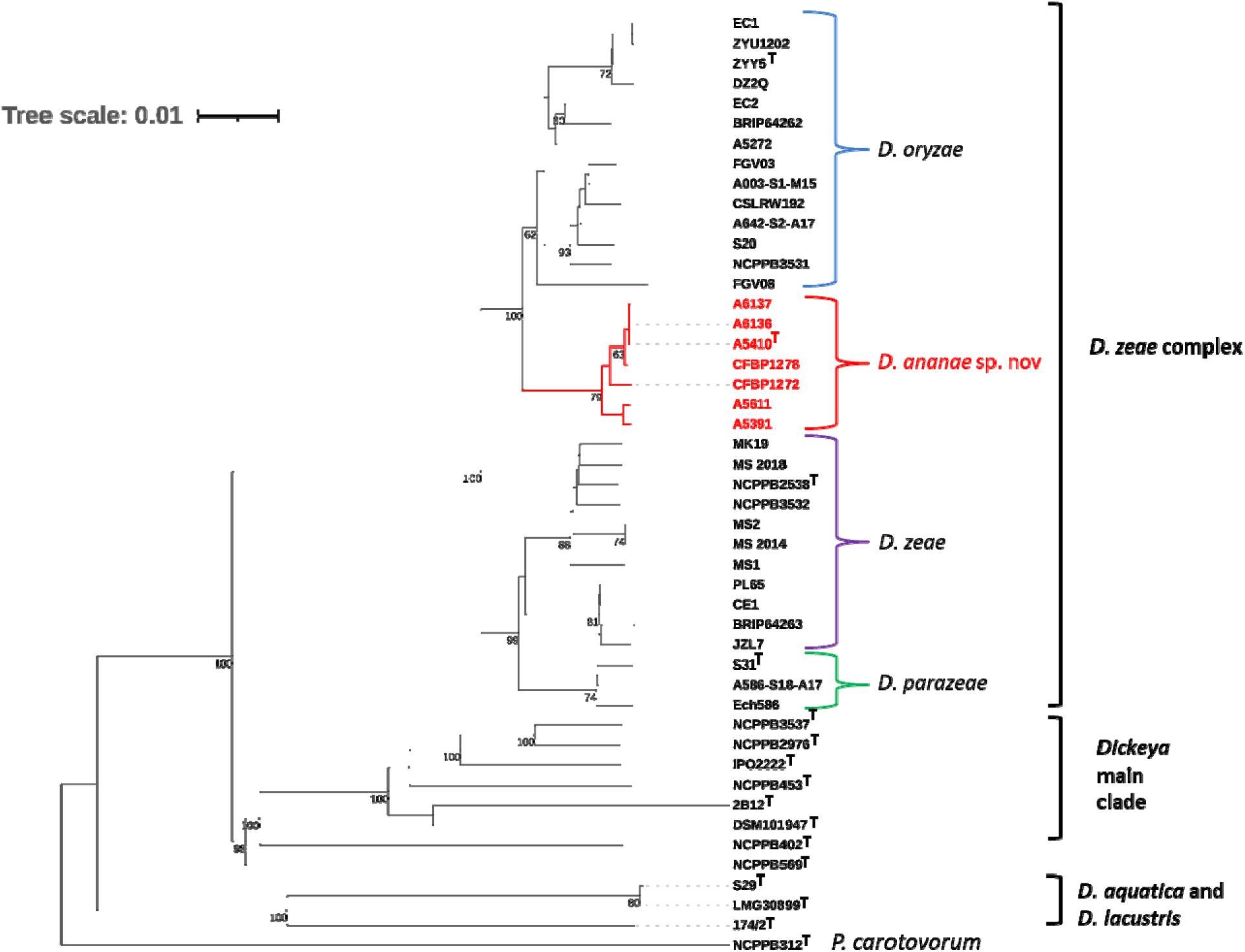
Whole genome sequence-based phylogenetic tree of the genus *Dickeya*. This tree wa constructed using the Type Strain Genome Server (TYGS), it shows the relationships between *D. ananae* sp. nov. A5410^T^ and strains of different *Dickeya* species. The tree was inferred with FastME 2.1.6.1 using the Genome BLAST Distance Phylogeny (GBDP) approach, with distances calculated from genome sequences via the TYGS server. Branch lengths are scaled in terms of GBDP distance formula d5. The numbers above the branches are GBDP pseudo-bootstrap support values >60% from 100 replications, with an average branch support of 92.7%. The tree was rooted at the midpoint (Farris, 1972). Leaf labels are annotated by affiliation to species and subspecies clusters, genomic G+C content, δ values and overall genome sequence length, number of proteins, and the kind of strain. The phylogenetic trees with bootstrapping value and colored annotations were created using web-based tool Interactive Tree of Life (iTOL v6).

A pan-genome dendrogram of the *D. ananae* sp. nov. strains, along with other *Dickeya* species, was constructed using Roary, based on the prevalence of different gene categories, including core genes (shared by 99–100% of the strains), soft core genes (shared by 95–99% of the strains), shell genes (shared by 15–95% of the strains), and cloud genes (shared by 0–15% of the strains). The strains in genus *Dickeya*, were found to contain 553 core, 73 soft core, 5,218 shell, and 19,068 cloud genes (Figure 7). The pan-genome dendrogram revealed that all seven strains of *D. ananae* were grouped into a single clade with a high bootstrap support value of 100%, showing significant species delineation and consistent placement of the strains, further corroborating previous phylogenetic analyses and supporting the proposal of *D. ananae* as a new species. The Roary matrix, derived from the analysis of 14,912 genes, indicates the diversity within the genus *Dickeya* (Figure 7). Additionally, A5410^T^ had 312 unique genes when compared to other type strains in the genus (Figure 8). The monophyly of *D. ananae* sp. nov. strains within the *D. zeae* complex are consistent and congruent across the phylogenetic and phylogenomic evaluations, with strong support from high bootstrap values, confirming the assignment of these strains as a novel species within the cluster.

**Figure 7.**
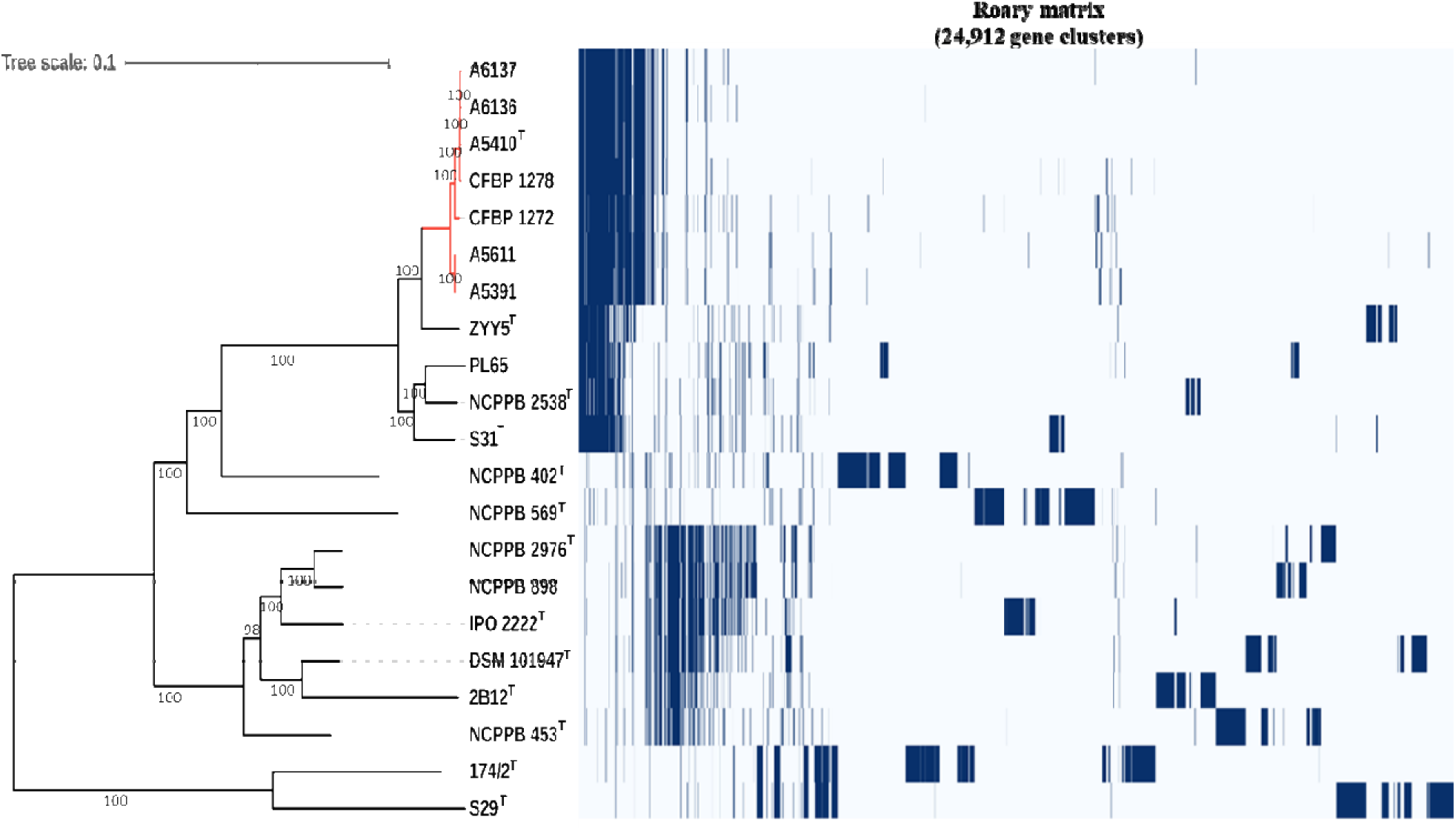
Core and pan genome analyses of *Dickeya* species including *D. ananae* sp. nov. The tree was constructed using 21 strains belonging to various *Dickeya* species. Left: Core-genome-based ML phylogenetic tree. Tree scale bar represents the nucleotide substitutions per site. Right: Pan genome-based Roary matrix of present and absent genes among the *Dickeya* species. The presence (blue) and absence (white) of accessory genome elements is shown.

**Figure 8.**
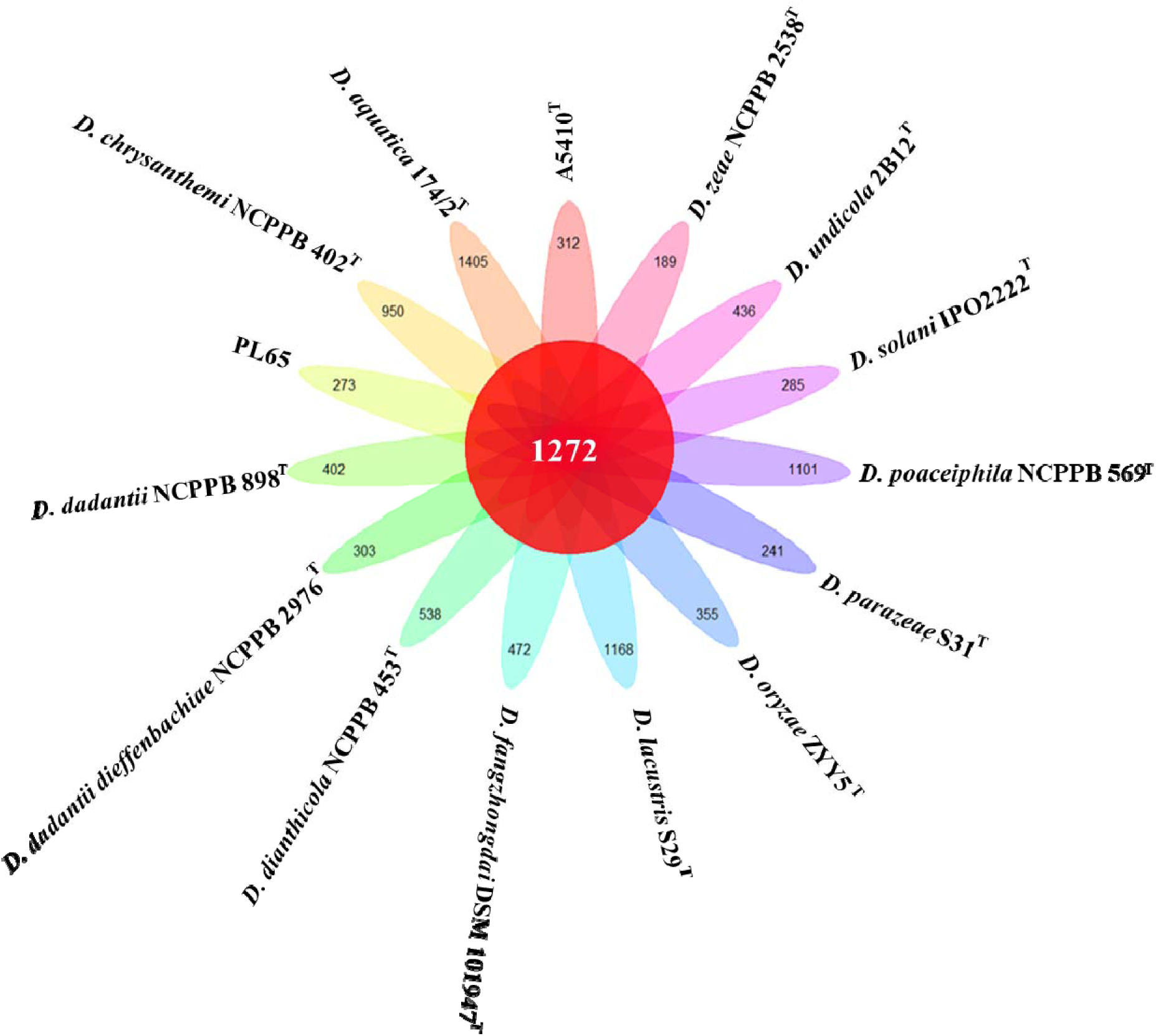
Pan-genome analyses of *Dickeya* species including *D. ananae* sp. nov. Pan-genome analyses of *D. ananae* sp. nov. (A5410^T^) with type strains of the other *Dickeya* species. Floral plot depicting the number of core orthologous genes (center) and the number of unique gene (petals). T in superscript denotes type strain of respective *Dickeya* species.

### Phenotypic, enzymatic and biochemical characterization

Electron microscopic investigations revealed that A5410^T^ exhibited a rod-shaped morphology. The cells of *D. ananae* sp. nov. measured approximately 2.91 ± 0.02 μm in length and 1 ± 0.07 μm in width, with peritrichous flagella (Figure 9), a characteristic trait observed in soft rot *Pectobacteriaceae* [66]. Motility assays using swimming and swarming methods showed that all strains exhibited swimming motility. The strains isolated from Hawaii demonstrated similar motility, with radii ranging from 5.35 ± 0.9 to 6.7 cm. However, the strains from Malaysia, CFBP 1278 (A5417) and CFBP 1272 (A5421), exhibited significantly weaker motility, with radii of 0.6 cm and 2.5 ± 0.56 cm, respectively, compared to the strains isolated from pineapples in Oahu. The macerating ability of these strains, previously demonstrated on pineapple, showed that all strains of *D. ananae* sp. nov. were capable of causing maceration of pineapple leaves [31]. All strains, including those isolated from Malaysia, were able to utilize pectate, as evidenced by pit formation on crystal violet pectate medium. Additional maceration tests on potato, taro, and onion slices revealed that all strains could macerate potato and taro, but not onion slices. Notably, all strains displayed stronger macerating activity on potato compared to taro slices (data not shown).

**Figure 9.**
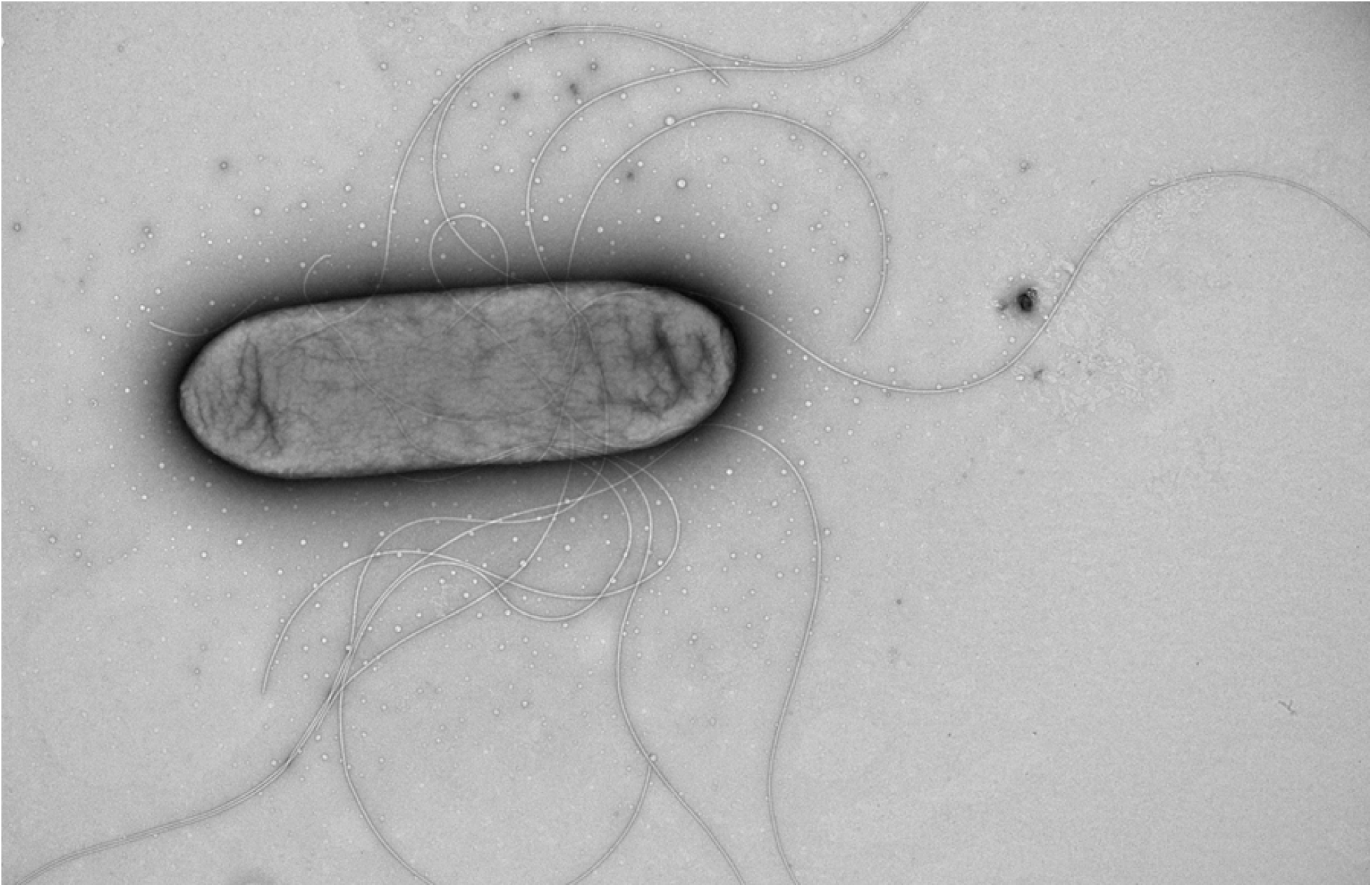
Transmission electron micrograph of *D. ananae* sp. nov. A5410^T^. A minimum of 10 images were captured for the analyzed strain across three biological replicates. A representative image is shown. Scale bar indicates 800 nm.

Using Biolog GENIII plates, the *D. ananae* sp. nov. members responded positively to the following carbon sources: sucrose, D-raffinose (with A5611, A5391, and CFBP 1272 showing weak positivity), D-melibiose, β-methyl-D-glucoside, D-salicin, N-acetyl-D-glucosamine, D-glucose, D-mannose, D-fructose, D-galactose, D-mannitol, and myo-inositol (except for CFBP 1278 and CFBP 1272) (Supplementary Table 4). Other positive responses included glycerol, D-glucose-6-PO_4_, D-fructose-6-PO_4_, D-aspartic acid (with CFBP 1278 weakly positive), L-aspartic acid, L-serine, pectin, D-galacturonic acid, L-galactonic acid lactone, D-gluconic acid, D-glucuronic acid, mucic acid, D-saccharic acid, L-lactic acid, citric acid, D-malic acid, L-malic acid, and bromo-succinic acid. All strains tolerated some inhibitory conditions, including pH 6, 1% NaCl, 1% sodium lactate, troleandomycin, rifamycin SV, lincomycin, Niaproof 4, vancomycin, tetrazolium violet, and tetrazolium blue. The strains showed inhibition to 4% NaCl, 8% NaCl, D-serine, troleandomycin, minocycline, nalidixic acid, potassium tellurite (except A5410^T^, which weakly tolerated it), lithium chloride (except A6136 and A5391, which weakly tolerated it), aztreonam, and sodium bromate. (Supplementary Table 4). The type strain A5410^T^ was unable to utilize the following carbon sources: D-serine, p-hydroxy-phenylacetic acid, D-lactic acid methyl ester, α-hydroxy-butyric acid, β-hydroxy-butyric acid, α-keto-butyric acid, and propionic acid. Some differences were noticed between the type strain A5410^T^ and the other type strains of the *D. zeae* cluster. In comparison to the strains *D. oryzae* NCPPB 3531^T^ and *D*.

*parazeae* S31^T^, A5410^T^ was able to weakly utilize D-lactose, N-acetyl-D-galactosamine, and inosine (Supplementary Table 4). Moreover, unlike the strains *D. oryzae* NCPPB 3531^T^, *D. parazeae* S31^T^, and *D. zeae* NCPPB 2538^T^, *D. ananae* A5410^T^ exhibited weak growth at pH 5 and was inhibited by troleandomycin (Supplementary Table 4).

The API 20E kit is specifically designed for identifying *Enterobacteriaceae* and other non-fastidious Gram-negative rods. It includes tests suitable for both genus-level identification and species characterization. The seven *D. ananae* strains were positive for indole production and acetoin production (Voges-Proskauer test), were able to utilize citrate, and exhibited β-galactosidase and gelatinase activity (Table 2). They were also able to ferment D-glucose, D-mannitol, L-rhamnose, D-saccharose, amygdalin, L-arabinose, and D-melibiose (except for A5391, which was unable to ferment D-melibiose). All strains were negative for the other tests, including H_2_S production, urease, arginine dihydrolase, lysine decarboxylase, ornithine decarboxylase (ODC), tryptophan deaminase. They were unable to ferment inositol and D-sorbitol. Nitrate reduction to nitrite was observed for all strains. The *D. ananae* strains sequenced in this study were negative for oxidase but were positive for gelatin hydrolysis, in contrast to *D. oryzae* strains, as reported by Wang *et al*. (2020) (Table 2).

**Table 2:** Biochemical and enzymatic characterization of tested strains in *D. ananae* sp. nov. using API 20E and API ZYM tests strips.

| Assays kit | Test | Strains |  |  |  |  |  |  |
| --- | --- | --- | --- | --- | --- | --- | --- | --- |
|  |  | A5410 (Type Strain) | A6136 | A6137 | A5611 | A5391 | A5417 (CFBP1278) | A5421 (CFBP1272) |
| API 20 E | β-Galactosidase (Ortho nitrophenyl-βD-galactopyranosidase, ONPG) | Yellow (+) | + | + | + | + | + | + |
|  | Arginine dihydrolase (ADH) | Yellow (-) | - | - | - | - | - | - |
|  | Lysine decarboxylase (LDC) | Yellow (-) | - | - | - | - | - | - |
|  | Ornithine decarboxylase (ODC) | Yellow (-) | - | - | - | - | - | - |
|  | Citrate utilization (CIT) | Blue (+) | + | + | + | + | + | + |
|  | H<sub>2</sub>S production (H<sub>2</sub>S) | colorless (-) | - | - | - | - | - | - |
|  | Urease (URE) | Yellow (-) | - | - | - | - | - | - |
|  | Tryptophan deaminase (TDA) | Brownish yellow (-) | - | - | - | - | - | - |
|  | Indole production (IND) | Pink (+) | + | + | + | + | + | + |
|  | Acetoin production (Voges Proskauer, VP) | Pink (+) | + | + | + | + | + | + |
|  | Gelatinase (GEL) | Diffusion of Black pigment (+) | + | + | + | + | + | + |
|  | D-Glucose (GLU) | yellowish green (+) | + | + | + | + | + | + |
|  | D-Mannitol (MAN) | Yellow (+) | + | + | + | + | + | + |
|  | Inositol (INO) | Blue (-) | - | - | - | - | - | - |
|  | D-Sorbitol (SOR) | Blue (-) | - | - | - | - | - | - |
|  | L-Rhamnose (RHA) | Yellow (+) | Blue green (-) | + | + | + | + | + |
|  | D-Saccharose (sucrose; SAC) | Yellow (+) | + | + | + | + | + | + |
|  | D-Melibiose (MEL) | Yellow (+) | + | + | + | Blue green (-) | + | + |
|  | Amygdalin (AMY) | Yellow (+) | + | + | + | + | + | + |
|  | L-Arabinose (ARA) | Yellow (+) | + | + | + | + | + | + |
|  | Nitrate Reduction (NIT) Reduction of Nitrates to No<sub>2</sub> | Red (+) | + | + | + | + | + | + |
| API ZYM | Alkaline phosphatase | Violet (+) | + | + | + | + | + | + |
|  | Esterase (C 4) | Violet (+) | + | + | + | + | + | + |
|  | Esterase (C 8) | Colorless (-) | - | - | - | - | - | - |
|  | Lipase (C14) | Colorless (-) | - | - | - | - | - | - |
|  | Leucine arylamidase | Orange (+) | + | + | + | + | + | + |
|  | Valine arylamidase | Colorless (-) | - | - | - | - | - | - |
|  | Cystine arylamidase | Colorless (-) | - | - | - | - | - | - |
|  | Trypsin | Colorless (-) | - | - | - | - | - | - |
| | α-chymotrypsin | Colorless (-) | - | - | - | - | - | - |
|  | Acid phosphatase | Violet (+) | + | + | + | + | + | + |
|  | Naphthol-AS-BI-phosphohydrolase | Blue (+) | + | + | + | + | + | + |
| | α-galactosidase | Colorless (-) | - | - | - | - | - | - |
| | β-galactosidase | Violet (+) | + | + | + | + | + | + |
| | β-glucuronidase | Colorless (-) | - | - | - | - | - | - |
| | α-glucosidase | Colorless (-) | - | - | - | - | - | - |
| | β-glucosidase | Violet (+) | + | + | + | + | + | + |
| | N-acetyl-β-glucosaminidase | Colorless (-) | - | - | - | - | - | - |
| | α-mannosidase | Colorless (-) | - | - | - | - | - | - |
| | α-fucosidase | Colorless (-) | - | - | - | - | - | - |
| **Other tests** | CVP (pit formation) | + | + | + | + | + | + | + |
|  | Oxidase (cytochrome-oxidase) | - | - | - | - | - | - | - |
|  | Catalase | + | + | + | + | + | + | + |

+, positive; - negative

Enzymatic reactions tested with the API ZYM kit revealed that *D. ananae* strains exhibit activity for the enzymes alkaline phosphatase, esterase (C4), leucine arylamidase, naphthol-AS-Bi-phosphohydrolase, β-galactosidase, and β-glucosidase. They were negative for esterase lipase (C8), valine arylamidase, cystine arylamidase, trypsin, α-chymotrypsin, α-galactosidase, β-glucuronidase, α-glucosidase, N-acetyl-β-glucosaminidase, α-mannosidase, and α-fucosidase (Table 2). The API 50 CH kit is designed for studying the metabolism of carbohydrates and carbohydrate derivatives, while the API 50 CHB/E medium is specifically designed for use with *Enterobacteriaceae*. All strains of *D. ananae* sp. nov. were able to produce acids from glycerol, D-arabinose, L-arabinose, D-ribose, D-xylose, D-galactose, D-glucose, D-fructose, D-mannose, L-rhamnose, D-mannitol, N-acetylglucosamine, arbutin, esculin ferric citrate, salicin, D-cellobiose, D-melibiose, D-sucrose. The seven strains showed a weak response for acid production with potassium gluconate. These strains showed variable responses in acid production from inositol (positive or weak), raffinose (positive or weak) and lactose (positive, weak, or negative) (Supplementary Table 5). All strains of *D. ananae* sp. nov were not able to produce acids from erythritol, L-xylose, D-adonitol, methyl-ß-D-xylopyranoside, L-sorbose, dulcitol, D-sorbitol, methyl-α-D-mannopyranoside, methyl-α-D-glucopyranoside, amygdalin, D-maltose, D-trehalose, inulin, D-melezitose, starch (amidon), glycogen, xylitol, gentiobiose, D-turanose, D-lyxose, D-tagatose, D-fucose, L-fucose, D-arabitol, L-arabitol, potassium 2-ketogluconate and potassium 5-ketogluconate. When compared to *D. oryzae* ZYY5^T^, differences were observed in the production of acids from D-galactose, D-glucose, D-fructose, D-mannitol, N-acetylglucosamine, and D-sucrose, as ZYY5^T^ was reported to be unable to produce acids from these carbon sources [26]. In contrast, strains of *D. ananae* sp. nov. were unable to produce acids from L-sorbose, unlike *D. oryzae* ZYY5^T^, which has been reported to produce acid from this carbohydrate [26]. Additionally, *D. ananae* sp. nov. could not produce acid from potassium 2-ketogluconate, similar to *D. oryzae* ZYY5^T^, but differing from the *D. zeae* type strain, which was capable of acid production from this carbohydrate [26].

Pectate lyases, cellulases, and proteases are important extracellular enzymes playing critical role in *Dickeya* pathogenesis. All strains of *D. ananae* sp. nov. possess pectinase activity, with A5410^T^, A6136, and A6137 exhibiting the highest levels of activity followed by A5391 and A5611. In contrast, strains CFBP 1278, and CFBP 1272 displayed weaker activity, as determined by semi-quantitative assays. Similarly, for cellulase activity, A5410^T^, A6136, and A6137 showed the highest activity, followed by A5391, A5611, CFBP 1272, and CFBP 1278, which exhibited the least activity. Regarding protease activity, *D. ananae* sp. nov. strains A5410^T^, A6136, A6137, A5391, and A5611 demonstrated comparable protease activity, while the Malaysian strains CFBP 1272 and CFBP 1278 showed only very weak activity after 30 hours of incubation (Supplementary Table 4).

Various degrees of sensitivity were observed across the six major classes of antibiotics tested, which included a total of 13 antibiotics (Supplementary Table 6). For the aminoglycoside class, the seven *D. ananae* strains exhibited similar responses, showing sensitivity to spectinomycin (100 mg/ml), gentamicin (50 mg/ml), and kanamycin (50 mg/ml). All strains showed equal sensitivity to chloramphenicol (50 mg/ml), trimethoprim (25 mg/ml), and carbenicillin (50 mg/ml). A5611 showed the highest sensitivity to vancomycin (50 mg/ml) with an inhibition zone of 1.85 cm. Differences in sensitivity to bacitracin (50 mg/ml), hygromycin B (50 mg/ml), and tetracycline (40 mg/ml) were noted among the strains In the β-lactam class, CFBP 1278 showed the highest sensitivity to penicillin (50 mg/ml) with a 2.15 cm inhibition zone, while A5410^T^ demonstrated the greatest sensitivity to cefalexin (50 mg/ml), with a 2.1 cm inhibition zone. All strains exhibited similar sensitivity to cefotaxime (50 mg/ml), with inhibition zones ranging from 2.4 to 3 cm.

In conclusion, based on the results of this polyphasic study, strain A5410^T^ is proposed to represent a novel species in the genus *Dickeya*, named *Dickeya ananae* sp. nov.

Description of *Dickeya ananae* sp. nov. (a.na.nae. L. F. gen. n. *ananas* isolated from pineapple)

*Dickeya ananae (*a.na.nae. L. F. gen. n. *ananas* referring to the isolation source of the type strain, pineapple).

*Dickeya ananae* is a Gram-negative, rod-shaped, motile bacterium measuring 2.91 ± 0.02 μm in length, with peritrichous flagella. It is fermentative and grows optimally at 28 °C on nutrient agar medium supplemented with 0.4% glucose and on King’s B medium, forming milky white colonies within 24 hours. This bacterium forms pits on crystal violet pectate medium within 24-48 hours [33] and causes maceration of potato tubers, taro slices, and pineapple leaves. *Dickeya ananae* sp. nov. is catalase positive, positive for nitrate reduction, oxidase negative and pectinolytic. *D. ananae* sp. nov. is able to utilize sucrose, D-raffinose, D. melibiose, β-methyl-D-glucoside, D-salicine, N-acetyl-D-glucosamine, D-glucose, D-mannose, D-fructose, D-galactose, D-sorbitol, D-mannitol, myo-inositol, glycerol, D-glucose-6-PO_4_, D-fructose-6-PO_4,_ D-aspartic acid, L-aspartic acid, L-glutamic acid, L-serine, pectin, D-galacturonic acid, L-galactonic acid lactone, D-gluconic acid, D-glucuronic acid, methyl pyruvate, L-lactic acid, citric acid, D-malic acid, L-malic acid, bromo-succinic acid, acetic acid and formic acid as a carbon sources. It shows inhibition at pH 5, or in the presence of 8 % NaCl, D-serine, minocycline, nalidixic acid, potassium tellurite, aztreonam and sodium bromate. The type strain is unable to utilize the following carbon sources: dextrin, stachyose, N-acetyl neuraminic acid, D-serine, quinic acid, p-hydroxy-phenylacetic acid, D-lactic acid methyl ester, tween 40,-amino-butryric acid, α-hydroxy-butyric acid, β-hydroxy-D, L-butyric acid, α-keto-butyric acid and propionic acid. Enzymatic reactions tested with the API ZYM kit give positive results for alkaline phosphatase, esterase, leucine arylamidase, acid phosphatase, naphthol-AS-BI-phosphohydrolase, β-galactosidase, and β-glucosidase. The type strain is able to utilize citrate, produces indole and acetoin and able to hydrolyze esculin; it is negative for urease. The API 50CH kit gave positive results for the utilization of glycerol, D-arabinose, L-arabinose, D-ribose, D-xylose, D-galactose, D-glucose, D-fructose, D-mannose, L-rhamnose, D-mannitol, *N*-acetylglucosamine, arbutin, aesculin ferric citrate, salicin, lactose, melibiose, sucrose, raffinose, and weakly positive for inositol and potassium gluconate. Assay for antibiotic susceptibility showed that *D. ananae* A5410^T^ is sensitive to 13 antibiotics including trimethoprim, kanamycin, bacitracin, cephalexin, hygromycin B, vancomycin, cefotaxime, tetracycline, spectinomycin, chloramphenicol, carbenecillin, penicillin and gentamicin. The highest sensitivity was observed for the cefotaxime, carbenicillin and cefalexins. The *D. ananae* type strain is A5410^T^ (=ICMP 25020^T^ =LMG 33197^T^) isolated from Pineapple in Hawaii. Its genome size of 4.77 Mbp with a G+C content of 53.5 mol% based on the genome sequence. Other isolates of this species were also obtained from pineapple in Hawaii (A5391, A6136, A6137, and A5611), and two strains, CFBP 1272 and CFBP 1278, were isolated from pineapple in Malaysia.

## Acknowledgements

The authors express their gratitude to Dr. Bevan Weir and Ms. Megan Petterson from ICMP, New Zealand, and Ms. Els Vercoutere and Ms. Claudine Vereecke from the BCCM/LMG bacteria collection for their assistance with strain registration in the respective culture collections. The Illumina library preparations and sequencing were carried out at the UC Davis Genome Center DNA Technologies and Expression Analysis Core, supported by NIH Shared Instrumentation Grant 1S10OD010786-01.

## Legends for supplementary Figures and Tables

**Supplementary Figure 1.** Maximum-likelihood phylogenetic tree based on 16S rRNA sequences comparison showing the position of *D. ananae* sp. nov. A5410^T^ within the genus *Dickeya.* The *Pectobacterium carotovorum* NCPPB 312 was taken as an outgroup. The phylogenomic tree based on 16S rRNA genome sequences in the TYGS tree inferred with FastME 2.1.6.1 (Meier-Kolthof and Göker 2019) from the Genome BLAST Distance Phylogeny approach (GBDP); distances were calculated from genome sequences. The phylogenetic trees with bootstrapping value and colored annotations were created using web-based tool Interactive Tree Of Life (iTOL v6). The branch lengths are scaled in terms of GBDP distance formula d5. The numbers above the branches are GBDP pseudo-bootstrap support values>60% from 100 replications. The tree was rooted at the midpoint.

**Supplementary Table 1.** Subsystem Annotation summary of 27 categories of subsystem features as predicted by RAST web server for new sp. A5410^T^ and the other genomes used in this study.

**Supplementary Table 2.** Combined Average nucleotide identity (ANI) (above diagonal) calculated using the OrthoANI algorithm implemented in the software OAT and digital DNA-DNA hybridization (dDDH) determined using Genome-to-Genome Distance Calculator version 3.0 (https://ggdc.dsmz.de/) (below diagonal) for all strains grouping in *D. ananae* sp. nov and other strains from NCBI database belonging to the *D. zeae* complex. Combined Average nucleotide identity (ANI) (above diagonal) and alignment percentage (AP) (below diagonal) calculated using CLC Genomics workbench 22.0.2 for all strains grouping in *D. ananae* sp. nov and other strains from NCBI database belonging to the *D. zeae* species complex. Da, *D. ananae*; Dor, *D. oryzae*; Dz, *D. zeae;* Dpz, *D. parazeae*.

**Supplementary Table 3.** List of *Dickeya* strains of and *Pectobacterium carotovorum* (outgroup) used in the *gapA,* 16S rRNA, MLSA, and whole genome phylogenetic analyses.

**Supplementary Table 4.** BIOLOG GEN III and other phenotypic profiles of *D. ananae* sp. nov. compared to *D. zeae*, *D. oryzae* and *D. parazeae*.

**Supplementary Table 5.** Fermentation of different carbon sugar by *D. ananae* sp. nov. determined using API 50 CH strips.

**Supplementary Table 6.** Antibiotic sensitivity assays of *D. ananae* sp. nov. against a panel of 13 different antibiotics.

